# VirStrain: a strain identification tool for RNA viruses

**DOI:** 10.1101/2020.12.21.423722

**Authors:** Herui Liao, Dehan Cai, Yanni Sun

## Abstract

Genome epidemiology, which uses genomic data to analyze the source and spread of infectious diseases, provides important information beyond interview-based methods. Given fast accumulation of sequenced viral genomes, a basic need in genome epidemiology is to identify which reference genomes are identical or closest to the ones in a sequenced sample. Then the associated metadata such as the geographical locations can be utilized to infer the transmission network. In this work, we deliver VirStrain, a fast and accurate tool for conducting strain-level analysis from short reads. By using a greedy covering algorithm, we are able to derive unique *k*-mer combinations for highly similar reference genomes. VirStrain is able to detect the most possible strain and also multiple strains that may simultaneously infect the same host. We tested VirStrain on three types of RNA viruses whose reference genomes have different similarity distributions. For each types of virus, we assessed VirStrain across multiple benchmark datasets of different properties and complexity. The experimental results on both simulated and real sequencing data show that VirStrain outperforms other strain identification tools.

## 2 Introduction

Advancements of sequencing technologies opened us new avenues for tracking the spread of infectious diseases. The most recent example is COVID-19. It triggered extensive efforts in sequencing SARS-CoV-2 from a large number of patients. By end of August 2020, there are 74,976 and 22,124 complete sequences publicly available from the Global Initiative on Sharing All Influenza Data (GISAID) [29] and GenBank [3], respectively. Besides SARS-CoV-2, there are also a large number of sequenced genomes for other RNA viral pathogens. For example, 152,938 and 927,873 genomes have been deposited into publicly available databases such as NCBI for Influenza and HIV infections. Compared to traditional methods that rely on incidence data and interview-based contact tracing, these genomes have provided important complementary information on analyzing the source, transmission, and spread of the diseases [7, 24, 11, 20, 10]. A fundamental step in applying genome epidemiology is to identify known strains that are identical or closest to those in a sequenced sample. Towards this goal, we will deliver a tool that can efficiently and accurately identify known strain for RNA viruses.

**Strain** As pointed out in [36], it is difficult to give a universal definition of microbial strain. Depending on the context, strain can refer to a variant of a given virus with unique and stable phenotypic characteristics under natural conditions [19], or a specific viral genome [36, 11, 15, 1, 12]. In this context, strain refers to a specific viral genome.

RNA viruses usually lack strict proofreading mechanisms during replication, leading to new copies containing genetic variations from the parent strains. Thus, it is not rare that genetic variations are found in genomes sequenced from different patients. Many of these variations can be neutral and deleterious to the virus survival. However, some mutations are beneficial to the fitness of the virus. For some extensively studied viruses such as HIV and Influenza, there are known mutations that lead to a different immune response in host cells, drug sensitivity and resistance [18, 25].

For newly discovered viruses such as SARS-CoV-2, whether or not the identified genetic changes affect the strains’ virulence and transmissibility needs substantial evidence. But the fast accumulation of genomic data sequenced from different patients worldwide has provided precious information for precision epidemiology for infectious disease control [20]. With associated metadata such as symptoms, geographical location, and travel history, the reference strain database can provide important information for inferring the transmission path when the contact history is ambiguous or missing.

Some studies that used the strain genomes for studying the spread of COVID-19 already showed that the clusters of the reference genomes are highly structured and are consistent with their geographical distributions [10]. Before COVID-19, analysis of strain genomes also made important contributions to precision epidemiology by local HIV breakout monitoring in Canada [26], Zika virus transmission study in Florida USA [9], and Ebola virus transmission in West Africa [23].

Given the development of NGS, sequencing the virus using either targeted or viral metagenomic sequencing methods from a large number of patients will become the norm during pandemics. Given short reads as input, we will output the reference genomes that are closest to the ones in the underlying sample. In addition, if subtype information is associated with the reference strain, users can also obtain the subtype, clade, or subclade of the virus.

The difficulty of strain identification depends a lot on the similarity between sequenced strains. Intensive sequencing efforts triggered by COVID-19 have resulted in a large number of genomes with high sequence similarity partially because of the recent association of this virus with human and also its relatively low mutation rate. Short reads tend to be ambiguously mapped or aligned to multiple reference genomes. Dissolving the ambiguity in the alignment is computationally expensive [1]. Faster methods based on genomic-specific *k*-mers often cannot reach the strain level resolution [33, 5]. In addition, the strain identification tool should be able to detect more than one reference strain if there are multi-strain infection, which is not rare for RNA viral diseases.

Although there are some strain-level analysis tools for bacteria, not all can be repurposed to take virus genomes as inputs. Some other tools’ cannot scale to the large number of strain genomes with high similarities. We summarize related work in the following section.

### 2.1 Related work

Popular taxonomic classification tools such as Kraken series [33, 4] are not able to conduct strain-level analysis when the reference genomes share high similarities. Our experimental results have shown that their performance on SARS-CoV-2 and Influenza is not satisfactory.

When near-complete virus genomes are available, alignment-based tools such as BLAST can be applied to find reference genomes with the highest similarity. If the inputs are short contigs or reads, alignment-based tools tend to align them to multiple genomes with the same scores. Similarly, other tools and websites that can monitor the mutations in strain genomes such as NextFlu and NextStrain [24, 11] also take genomes as input and assign the genome into major genetic groups. When there are difficulties in constructing high-quality virus genomes due to complexity of the data (e.g. metagenomic data), the low abundance of the virus, or the presence of a minor strain besides a major one, there is a need for a tool that can still identify the strains using reads as input.

There are some methods that focus on conducting strain-level analysis using short reads as input. Automated strain-level epidemiology analysis tools mainly focused on bacteria in microbiome sequencing data. The available reference-based strain-level analysis tools were divided into four groups by Yan et al. [36]. The first group focused on identifying known genotypes from reference genomes [13, 1]. The second group will further identify potentially novel strain of one bacterial species [31]. The third group of tools can identify one or multiple strains using reference-based methods [21, 27]. The last group uses structural variants rather than just single nucleotide variations (SNVs) [28], which usually renders better accuracy.

As the main goal of this work is to develop a method that can identify the closest strain to a known strain from short reads data, it is more related to the first group of the strain-level analysis tools for bacteria. The representative tools in the first group, PathoScope [13] and Sigma [1], rely on ambiguity-resolved read mapping strategies between short reads and reference genomes with high sequence similarity. Both tools allow users to create their own reference database and thus can be applied to viruses. However, they are too slow for identifying strains with tens of thousands reference genomes and large-scale sequencing data.

Other bacteria-centered tools cannot be conveniently re-purposed for virus strain analysis because they use bacteria-specific features such as bacterial marker genes, structural variants etc. For example, our experiments showed that existing marker gene sets can recognize HIV if the reads are sequenced from the dominant strain HXB2, but not other strains of HIV tarvir. One possible reason is that the derivation of the marker genes by existing programs [31] did not use all the available viral strains and thus can miss many strains.

In addition, RNA viruses usually possess higher mutation rates than bacteria because of the error-prone replication process. As a result, different patients contain different viral genomes. In particular, for a new breakout such as COVID-19, a large number of sequenced strains within a short period often share extremely high sequence similarity, making identifying individual strains from new data more challenging. Use the available haplotypes or strains to infer transmission has been applied to COVID-19. For example, Gudbjartsson et al. [10] are able to assign a sample to its closest haplotype based on a manually derived haplotype table. However, it is not clear the manually created table can scale to larger datasets or other viruses.

A very relevant work with the similar challenge is QuantTB [2], which targeted at identifying individual *M. tuberculosis* strains with high similarity. However, their tool is “hard coded” for *M. tuberculosis* and thus we cannot conveniently extend it to viruses. In addition, they also applied different thresholds on the number of distinct SNPs between strains, which are actually still stringent for newly identified RNA viruses such as SARS-CoV-2

## 3 Methods

In this work, we developed a tool, VirStrain, which can quickly identify one or multiple reference strains closest to those in short-read sequencing data. It achieves a better trade-off between speed and resolution by deriving unique *k*-mer combinations that can distinguish highly similar strains.

VirStrain conducts strain identification using short reads as input and doesn’t rely on sequence assembly, making it more amenable to cases where full virus genome cannot be assembled. The output of VirStrain contains the most possible strain (the strain that best matches the SNVs found in the sample set) identified in the data and the detailed read coverage of its single nucleotide variation (SNV) sites in an interactive HTML format (Supplementary Figure S7).

Highly similar reference genomes may not possess genome-specific *k*-mers. But they can possess genome-specific *k*-mer set, where the component *k*-mers can be utilized together to distinguish different reference genomes. Figure 1 shows a toy example of using *k*-mer sets to distinguish five sequences when there are no genome-specific *k*-mers. In order to find such *k*-mer set, we develop a greedy covering algorithm to identify unique combinations of SNV sites from aligned virus reference genomes. Then *k*-mers will be extracted from the SNV sites and construct *k*-mer set for underlying genomes.

**Figure 1:**
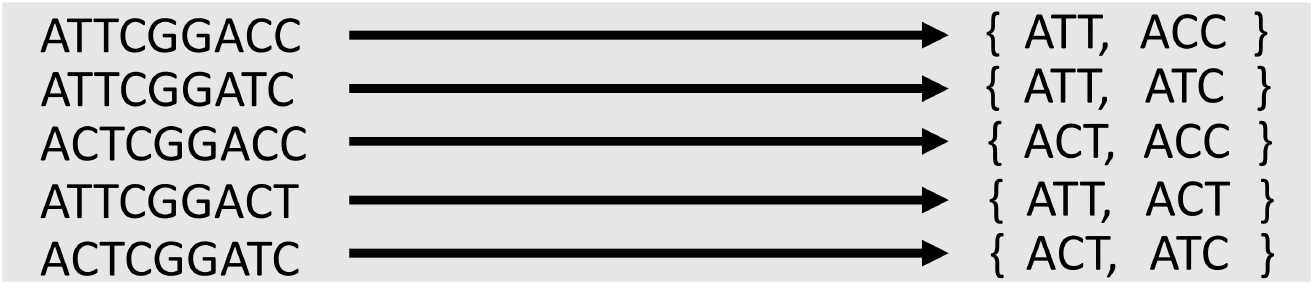
Use *k*-mer sets to distinguish five sequences of high sequence similarity. Each sequence has a unique *k*-mer combination.

### 3.1 Step 1: identify unique set of SNVs from reference genomes

The input to this algorithm is an MSA of the reference genomes. It is noteworthy that generating MSA for thousands to tens of thousands of genomes can be slow. But when the reference genomes share high sequence similarity (such as for SARS-CoV-2), the MSA can be produced using more efficient programs, such as the one provided by Mafft at its website [16].

Given the MSA M, the program will exclude all the sites where no SNV is observed. Instead, the algorithm favors variations from conserved sites, which indicate features that are specific to one or a small number of genomes. Thus, given M, we compute the Shannon entropy ℋ for each column.

Then we pick the column with minimum positive ℋ from M. Let this column be *s*_*i*_. Let the nucleotide at *s*_*i*_ with the minimum frequency be 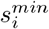. All the genomes containing 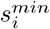 at site *s*_*i*_ will be extracted and saved in one cluster. The entropy for the remaining genomes will be updated after the extraction. And this greedy choice will be applied to the remaining genomes until all the genomes are in one cluster. It should be noted that low-quality columns with too many dashes will not be considered and can be filtered in pre-processing. Depending on the reference genome similarity and alignment quality, users can choose a threshold for the allowed percentage of dashes in one column. For SARS-CoV-2 and H1N1(HA), our default cutoff is 0. For HIV, our default cutoff is 10%. The pseudocode of the entropy-based greedy covering algorithm is presented in Algorithm 1. A working example is shown in the top-left panel of Figure 2.

**Figure 2:**
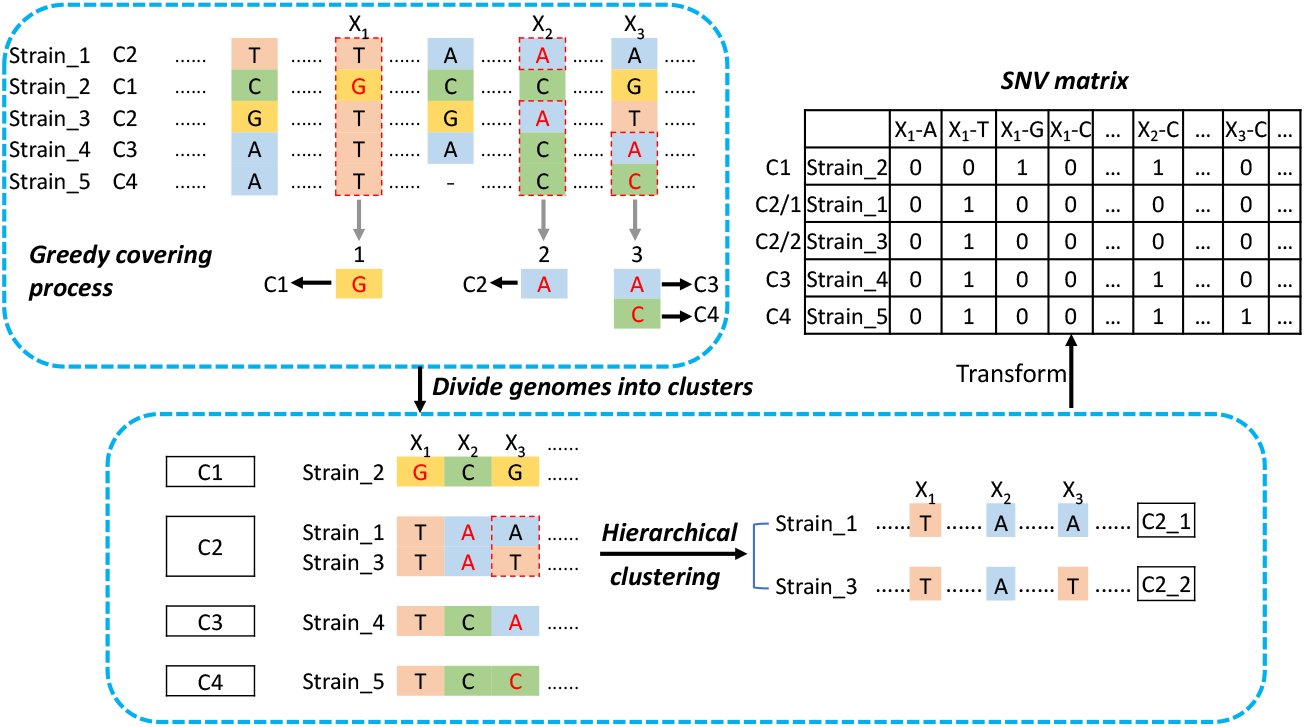
The sketch of the *k*-mer identification and SNV matrix construction stages. The SNVs in the matrix are represented by the site and base [2]. “C1-C5” represents the name of clusters.

After we apply this greedy covering algorithm, the reference genomes are divided into multiple clusters, where each cluster is defined by one SNV event. Figure 3 sketches the SNV sites for different clusters based on the order of SNV site selection in the greedy covering algorithm. Let the number of chosen SNV sites (i.e. the final number of clusters) be *m*. Let *s*_*i*_ be the SNV site chosen at the *ith* step in the greedy covering algorithm. Again, 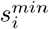 is the base with minimum frequency at site *s*_*i*_ in the remaining genomes. Let the corresponding cluster be *c*_*i*_, which contains the genomes containing 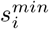 at the *ith* step. We use 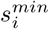 at SNV sites *s*_1_, *s*_2_, …, *s*_*i*_ to represent cluster *c*_*i*_. We have the following theorem and proof.

**Figure 3:**
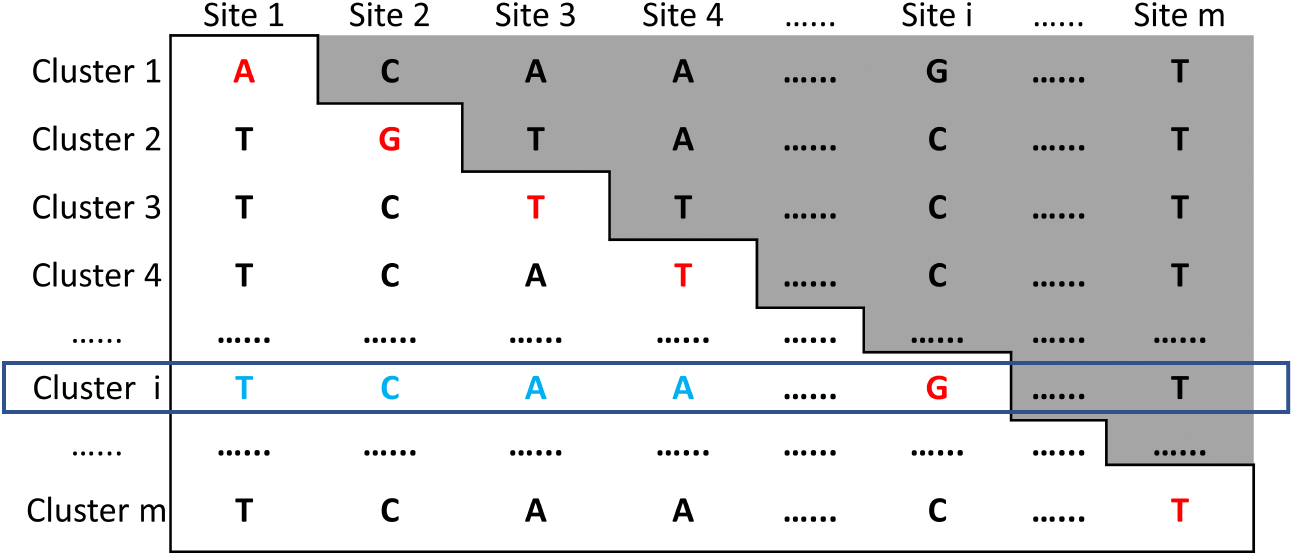
The SNV sites and corresponding clusters. Site *i* is the SNV site chosen at the *i*th step. The base in red denotes the base with the minimum *p*^*b*^ (i.e.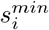). Genomes in the same cluster share the red base at the chosen SNV site. Each cluster has unique SNV base combination, which is shown in the white part of each row. The SNV base combination for cluster *c*_*i*_ is shown in a box.

#### Algorithm 1 Divide reference genomes into clusters with unique SNV combinations using the greedy covering algorithm

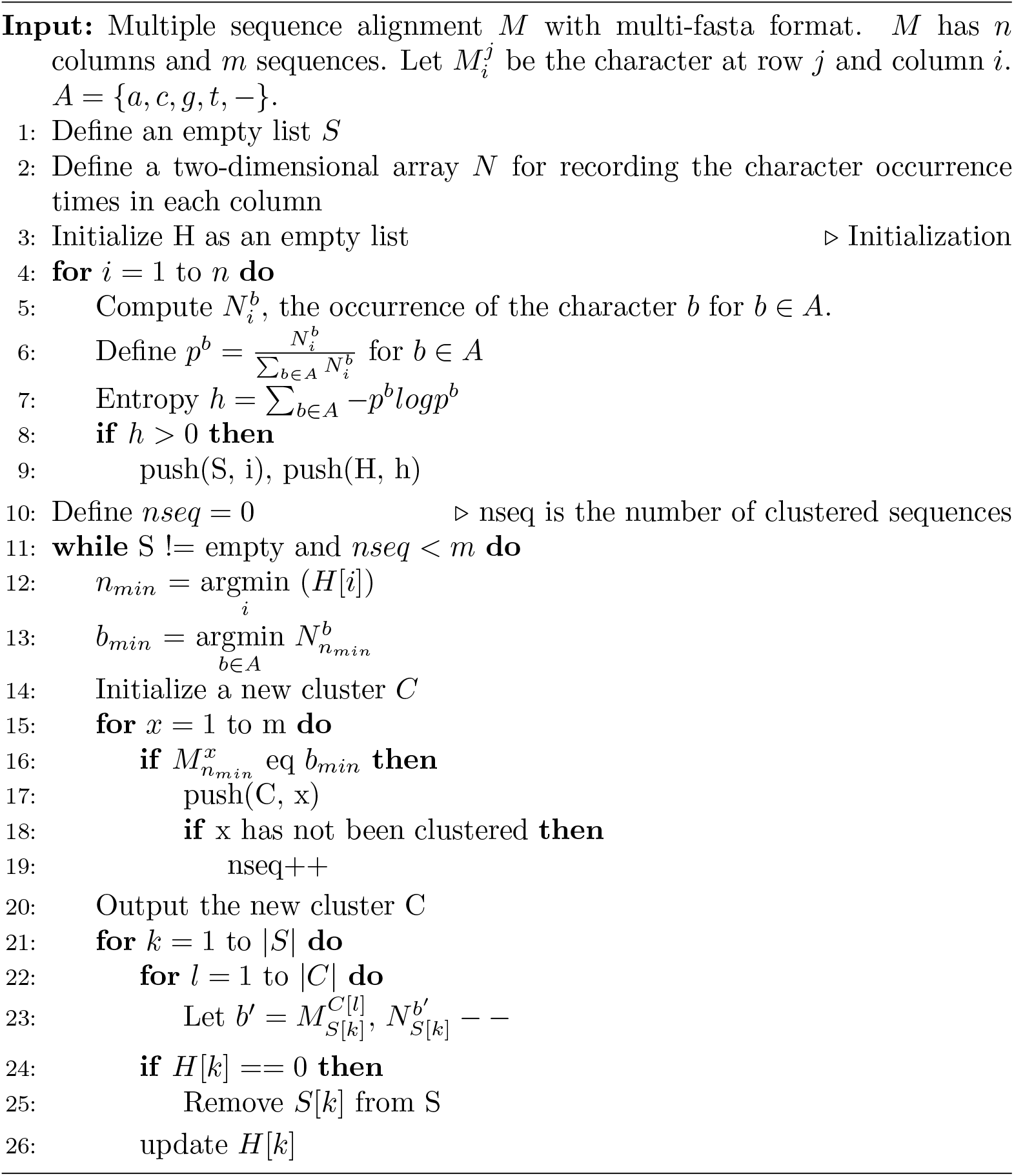

#### Theorem 1.

*Let the SNV base combination for the ith cluster c*_*i*_ *be* 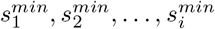. *This nucleotide base combination uniquely represent the genomes in cluster c*_*i*_.

*Proof*. In order to prove that the nucleotide base combination 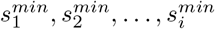 uniquely represent the genomes in cluster *c*_*i*_, we need to show that at least one base of these SNV events in *c*_*i*_ is different from any other cluster *c*_*j*_, where *j* ≠ *i*. Figure 3 can be used to illustrate this proof.

Without losing generality, we consider two cases. In case 1, we consider a cluster *c*_*j*_ with *j < i*. The bases at *j* in genomes of *c*_*j*_ must be identical (condition for clustering). In addition, that base must be different from the base at site *s*_*j*_ in genomes of cluster *c*_*i*_. Otherwise, the genomes of *c*_*i*_ will be clustered into *c*_*j*_ at step *j*. In the second case, we consider a cluster *c*_*j*_ with *j > i*. For any genomes in *c*_*j*_, their base at SNV site *s*_*i*_ must be different from the base at site 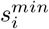 of the genomes in *c*_*i*_. Otherwise, that genome will be clustered into *c*_*i*_. Thus, we proved that at least one base at the SNV event combination in *c*_*i*_ is different from any other cluster *c*_*j*_.

Thus, as shown by Figure 3, we identified unique SNV combinations for each cluster. As we found the columns involving indels may have alignment errors or assembly errors (an example can be found in Supplementary Figure S6), we only use columns with no gap or a small number of gaps in greedy covering. As a result, some clusters can contain multiple genomes. In this case, the genomes inside each cluster can be aligned again (to reduce the alignment errors) and be clustered in a hierarchical fashion. An example can be found in the bottom panel of Figure 2.

#### 3.1.1 Maintaining “balanced” SNV site combinations

In the ideal case of no sequencing errors and each base of a viral genome being covered by at least one read, the SNV site sets that uniquely represent genomes in each cluster *c*_*i*_ are sufficient to determine the cluster or the strain precisely. But in reality, both heterogeneous coverage and sequencing errors exist. For clusters represented by a small number of SNV sites (e.g. *c*_1_ contains just one SNV site), sequencing errors can incur false positives. To address this issue, we will balance the number of the SNV sites for each cluster so that each cluster has the same number of SNV sites. To do so, we will use all *m* SNV sites for strain identification. If the original SNV sites can distinguish the genomes in different clusters, adding SNV sites will not change this property. Thus, each cluster still possesses unique SNV site combination. As shown in Figure 3, the SNV bases in both the white and gray part will be used for strain identification.

### 3.2 Step 2: iterative strain search algorithm

#### *k*-mer extraction

We will extract *k*-mers from these SNV sites, with the center base of each *k*-mer coming from this site (see Supplementary Methods 1.1). Figure S1 in the supplementary material shows an example of *k*-mer extraction. In order to avoid using *k*-mers that repeat at different sites in the MSA, we determine the *k*-mer size by examining the repeat times of *k*-mers of different *k* in the MSA. We find that with the increase of *k*, the repeat numbers of the *k*-mer at different sites reduce quickly (Supplementary Figure S2). By default, we use *25*-mer.

To detect all possible strains in a sample, we take an iterative strategy similar to QuantTB [2]. The overall workflow of the strain search algorithm is displayed in Figure 4. For an input set of reads, the *k*-mer match frequencies are computed using a *k*-mer counting tool and are mapped to an SNV matrix, which will allow us to quickly compute the sum of the coverage for all the SNV sites and rank the strains. The major operations are described below.

**Figure 4:**
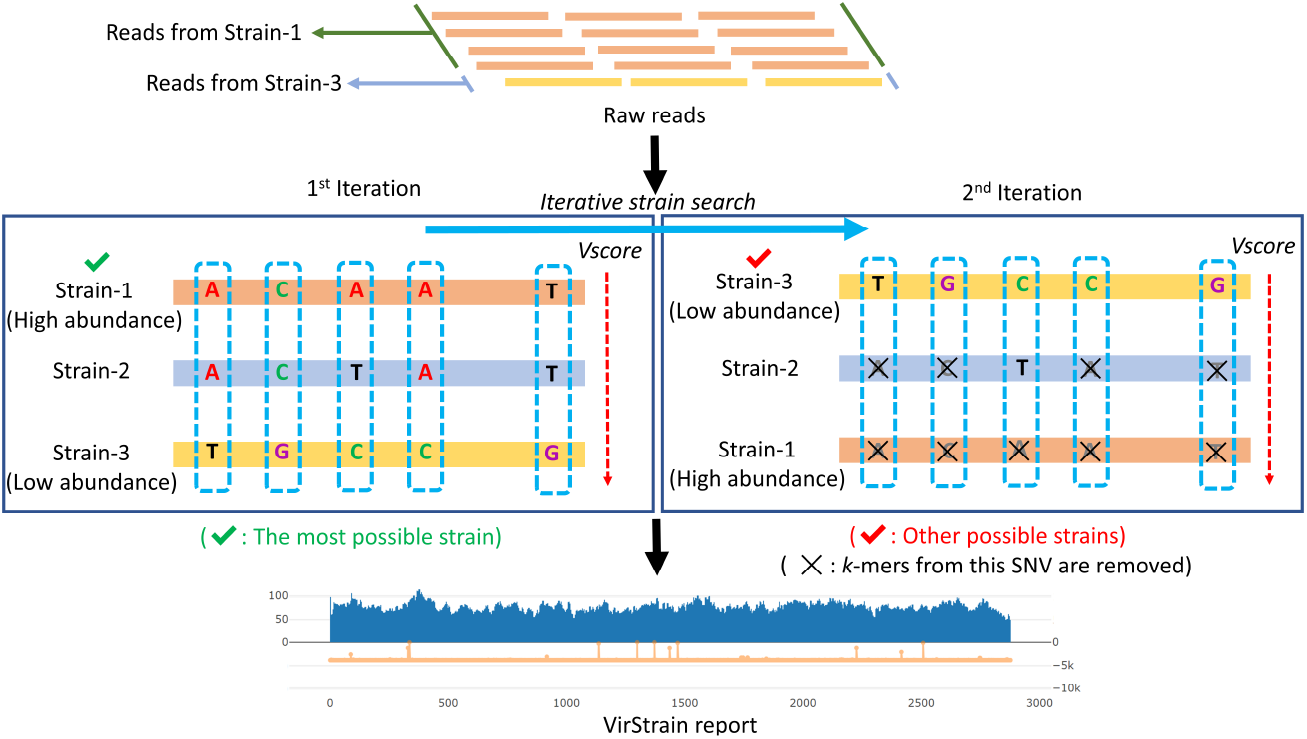
The overall workflow of the iterative strain identification. The red arrow represents that all strains are listed in descending order of *V score*.

#### Construction of the SNV matrix

The SNV sites chosen by the greedy covering algorithm will be used to construct an SNV matrix *S* of size 4*mn*, where *m* is the number of chosen variation sites and *n* is the number of reference genomes. An example is given in Figure 2. For a strain *i* and a chosen SNV site *x*, there are four cells corresponding to bases A, C, G, and T in *S*. Denote an SNV event as *x*-*b*, indicating that base *b* is observed at site *x*. A cell *S*_*i,x*−*b*_ is 1 if the strain *i* has base *b* at site *x*. Otherwise, it is 0. Each cell in the matrix *S* has associated *k*-mers match frequency.

#### Rank the reference genomes using *k*-mer match frequency

We apply Jellyfish (V2.3.0) [22], a fast multi-threaded *k*-mer counter, to count *k*-mers in the sequencing data. Let *F*_*x*−*b*_ be the *k*-mer match numbers of base *b* at site *x*. Thus, *S*_*i,x*−*b*_ = *S*_*i,x*−*b*_ * *F*_*x*−*b*_. Then, we will compute the frequency of base *b* at site *x* by normalize the *F*_*x*−*b*_. Therefore, 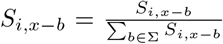. To reduce the effect of sequencing error, we filter *S*_*i,x*−*b*_ if its value is smaller than a given threshold.

Once *S* is updated based on the actual *k*-mer match frequency from the reads, we will compute the score of strain *i* using 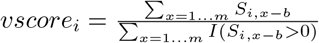, where *I* is an identity function. *vscore* favors strains with the most number of *k*-mer hits. Although it looks reasonable to consider other factors such as uniformity of *k*-mer match frequency, our empirical studies show that considering the total number of *k*-mer hits renders the best accuracy. One possible reason is the heterogeneous coverage of real sequencing data along RNA viral genomes. Read coverage profiles of 11 real sequencing datasets in our experiment can be found in Supplementary Figure S8. We will compute *vscore*_*i*_ for all the strains and rank them in decreasing order.

#### Iterative strain search

VirStrain takes an iterative approach to search for multiple strains. VirStrain will output the top 1 strain in the ranked list and then update *S* by replacing the frequency of all the SNV sites in identified strain with 0. Any strains that share the same SNV bases with the identified strains cannot reuse the frequency. Otherwise, strains that share high similarity with the identified ones can easily get higher *V score* than low-abundance strains that are not similar to the best match. An example is given in Figure 4. At each iteration, the sequencing depth is calculated by taking the average frequency of its SNVs for each identified strain in the sample. VirStrain continues to calculate the score and identifies the best matched strain in each iteration until the frequency values of all variations become 0. At each iteration, the sequencing coverage of the identified strain is calculated by taking the average of the *k*-mer match frequencies. In the end, this iterative process will return a list of strains with their *k*-mer coverage profiles on the SNV sites.

We sacrifice the resolution of finding highly similar strains in the same sample by avoiding introducing false positive hits via the iterative search strategy. If there are indeed highly similar strains such as those in quasispecies, the most abundant one will be output as a representative. We conducted experiments to examine how many different SNV sites are needed for VirStrain to recognize multiple strains (see Supplmentary Section 2.5).

## 4 Results

In order to test our tool on viruses with a large number of reference strain genomes and different mutations rates, we assessed our tool on three types of RNA viruses. The first is SARS-CoV-2, many of which have very high sequence similarity and may differ only at a few sites. The second is the “HA” region of Influenza A H1N1, which has lower average similarity than SARS-CoV-2 but higher similarity within the same clades and sub-clades. The third is HIV, which has a much lower similarity than SARS-CoV-2 and H1N1. For HIV, we used the “Gag” region, which is one of the marker genes for HIV subtype classification [30]. Figure 5A compares the pairwise Hamming distance at the SNV sites chosen by our algorithm from the aligned reference sequences.

**Figure 5:**
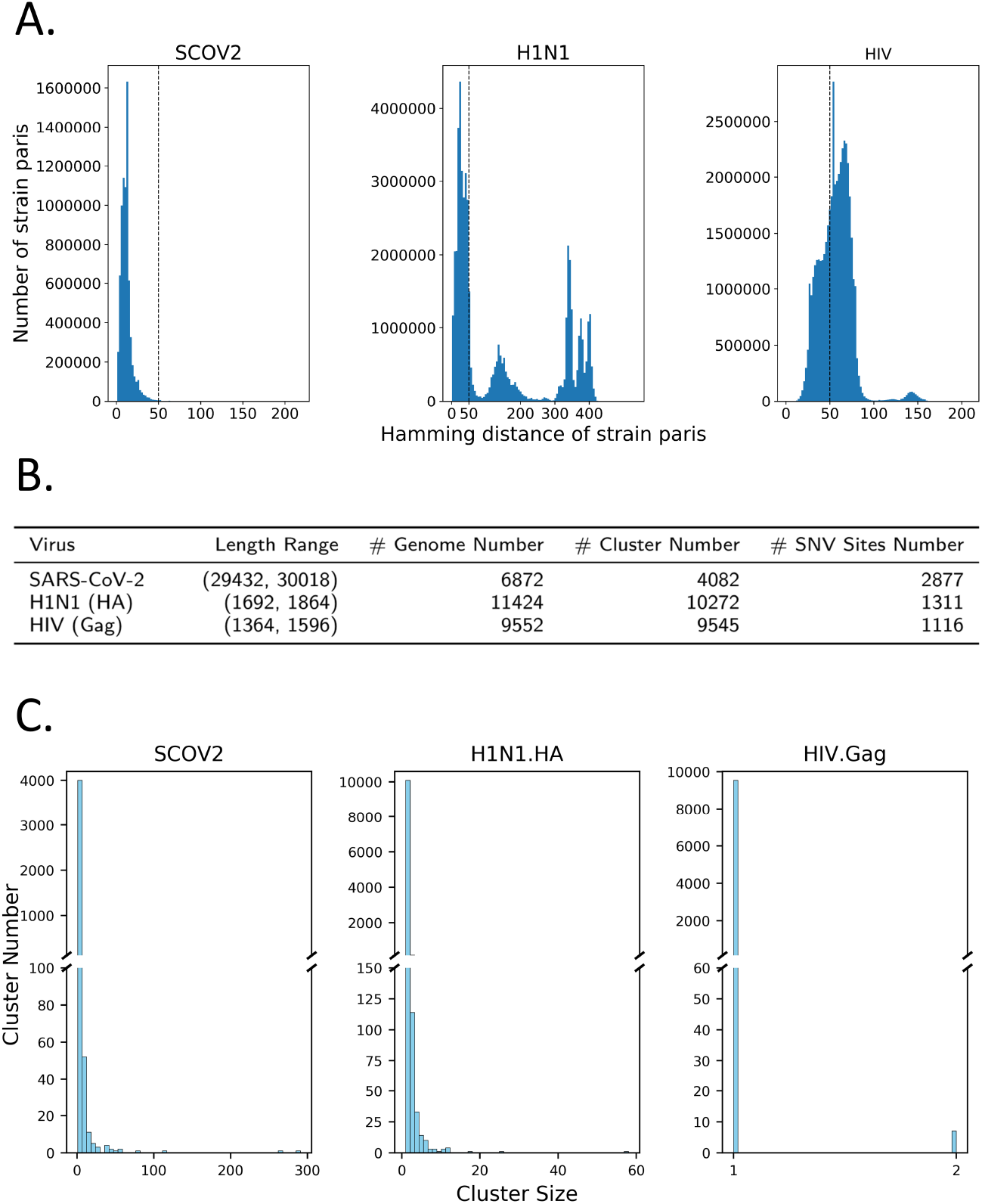
**(A)**. The pairwise Hamming distance distribution of SARS-CoV-2, H1N1 (HA), and HIV (Gag). The Hamming distance here is measured only on the chosen SNV sites by the greedy covering algorithm. They can represent the overall similarity of the three reference sets. The dashed line is at distance value 50 across the three panels for easy comparison. **(B)**. The statistics of the reference database of 3 viruses. “# Genome Number”: the number of downloaded genomes. “# Cluster Number”: the number of final clusters generated by VirStrain. “# SNV Sites Number”: the number of SNV sites chosen by VirStrain for each virus. **(C)**. The cluster size distribution of SARS-CoV-2, H1N1 (HA), and HIV (Gag). “Cluster size” represents the number of strains contained in each cluster.

### 4.1 Data and clustering results

To construct the reference database for VirStrain, we collected all available complete genomes of SARS-CoV-2 from NCBI, H1N1 (HA) from Influenza Research Database (IRD, http://www.fludb.org), and HIV (Gag) from HIV database (http://www.hiv.lanl.gov) as of July 14, 2020. Figure 5B summarizes the statistics of the reference genomes/genes of these three viruses, their final clustering results and also the number of their SNV sites chosen by VirStrain. It shows that the number of HIV (Gag) sequences is nearly equal to the number of clusters while the other 2 viruses differ a lot, which is caused by relatively low sequence similarity between HIV (Gag). The cluster size distribution of three viruses are displayed in Figure 5C. Most clusters are very small, with many containing a single genome. But there are also genomes that cannot be distinguished by the chosen SNV sites. Especially, there are strains with different lengths, leading to alignments with many gaps at the beginning and ending parts. Those columns are not utilized by the greedy covering algorithm. Thus, these strains are usually in the same cluster.

### 4.2 Overview of the experiments

The input to VirStrain are short reads from either relatively pure or highly-mixed samples (such as viral metagenomic data). VirStrain is able to directly return strains from both.

We assessed VirStrain from multiple aspects. The organization of all experiments is summarized in Figure 6. First, we focused on evaluating the possible limitations of VirStrain based on the method design (Figure 6A). In particular, we will evaluate how read length and sequencing coverage can affect the performance of VirStrain. Then we will investigate the applicability of VirStrain to mixed samples by conducting *k*-mer match against reads from the host, bacteria, etc. In addition, the iterative strain search poses a tradeoff between the resolution and accuracy of multi-strain identification. We will provide some guidance for users. Second, we benchmarked VirStrain against other popular strain-level analysis tools with simulated data (Figure 6B). Third, we validated VirStrain in multiple usage scenarios with both real data and mock data (Figure 6C).

**Figure 6:**
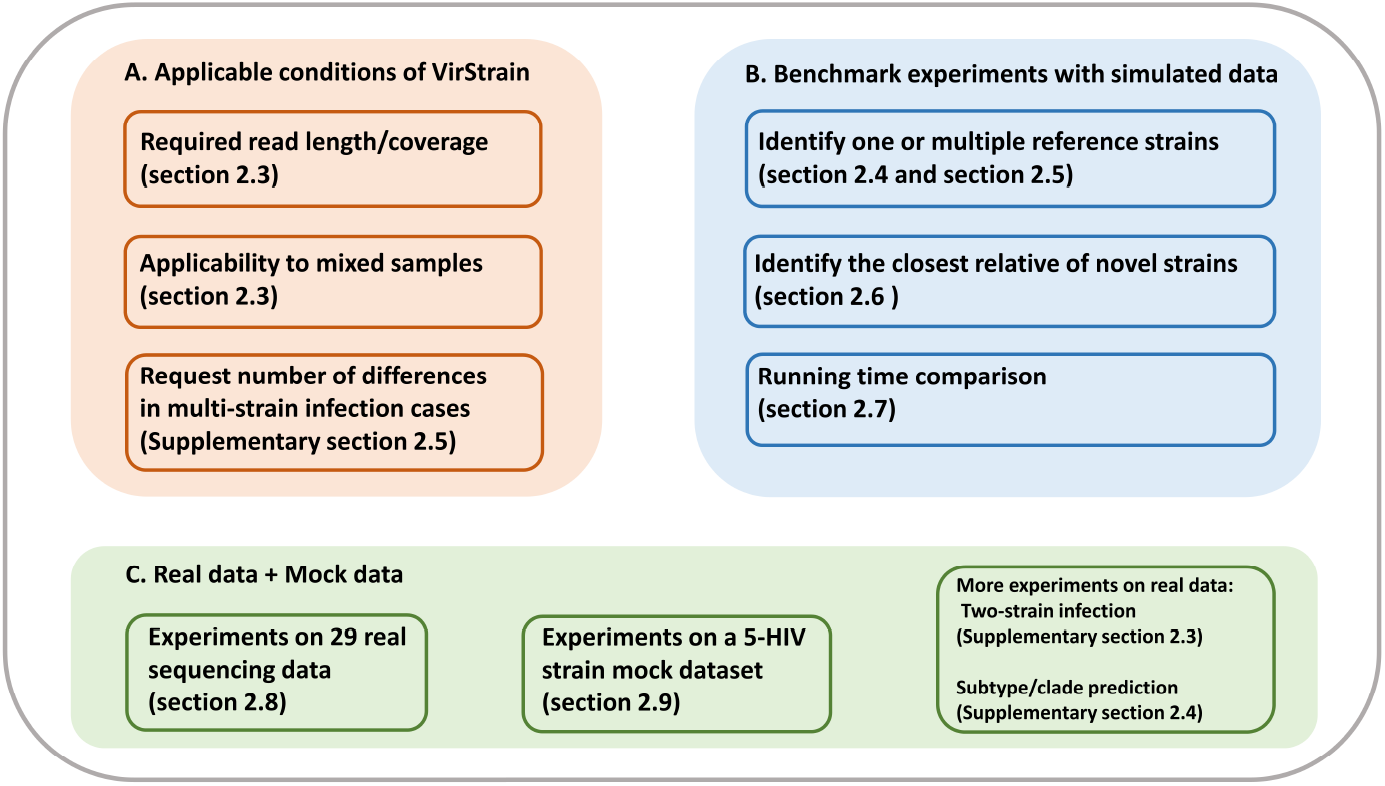
The overview of all experiments.

In all these experiments, we use accuracy as the main performance metric for different tools. It is worth noting that all tested tools usually output multiple strains with associated ranking. If we know the the number of strains (e.g. x) in a sample, we will keep only the top x outputs of a tool. Then the **accuracy** is defined as the percentage of correctly identified strains in the output. It’s noted that if multiple tied best matches are presented, with the correct strain among them, this will be counted as correct. We will quantify the number of “tie cases” in our experiments.

### 4.3 Baseline performance of VirStrain

The strain identification performance can be affected by strain coverage, read length, similarity of strains in the same sample. In order to provide users with guidance on utility of VirStrain, we conducted experiments to evaluate the robustness of VirStrain when the input data have different properties. First, we evaluated how the read length and strain coverage affect the performance of VirStrain. Second, we evaluated whether VirStrain could falsely identify other microbes as viruses, which is important for using VirStrain on highly mixed samples such as viral metagenomic data. Third, we focused on evaluating the minimum number of differences between the strains for VirStrain to identify them in the multi-strain infection case (Supplementary Section 2.5).

Because SARS-CoV-2 is of high interest, has large size, and high strain-level similarity, we conducted all experiments in this section using all our strains in SARS-CoV-2 that do not contain non-ACGT characters. So, there are 2,280 strains in the reference database.

#### Read length/coverage experiment

In order to evaluate the impact of read length and sequencing depth on VirStrain, we simulated reads from each single strain with 5 different sequencing depths and 4 different read lengths. Thus, there are altogether 20 combinations as shown in Table 1. For each combination, we conducted 2,280 experiments using ART simulated reads [14] from each of the 2,280 strain as input respectively. For all these experiments with different inputs, we found that the known reference strain always has the best *V score* (see Section 3.2). However, when the reads are too short and the coverage is low, there are many “tie cases” where multiple strains have the same *V score* as the reference strain. Table 1 shows the number of the tie cases out of the 2,280 experiments for each case and also the median number of strains in the top ranking group based on *V score*. For example, when the reads have the length 75bp and coverage 5x, 903 strains have the top ranking group with at least 2 strains. The median number of strains in this group is 7. With the increase of the coverage, the tie cases drop significantly. When the coverage is 10x, the median number of strains in the top ranking group is 2. With the increase of the coverage, the tie cases reduce to 0 and the top 1 strain is always the correct reference. Thus VirStrain should be applied to identify strains with at least 10x coverage. Otherwise, multiple strains of the same top *V score* will be returned. The change of read length does not significantly influence the performance when the depth is above 20x.

**Table 1:** The number of tie cases and the median number of best matches in all tie cases. Each cell contains a tuple with the first number being the number of tie cases and the right number being the median number of strains in the top-ranking group.

|  | 75bp | 100bp | 150bp | 250bp |
| --- | --- | --- | --- | --- |
| 5X | (903, 7) | (782, 6) | (602, 5) | (679, 5) |
| 10X | (141, 2) | (109, 2) | (51, 2) | (65, 2) |
| 20X | (0, 1) | (0, 1) | (0, 1) | (0, 1) |
| 50X | (0, 1) | (0, 1) | (0, 1) | (0, 1) |
| 100X | (0, 1) | (0, 1) | (0, 1) | (0, 1) |

#### Will the *k*-mers derived by VirStrain match non-viral genomes?

As highly mixed samples such as viral metagenomic data can contain reads from non-viruses, it is fair to ask whether VirStrain may construct false strains from non-viral reads. In order to evaluate this, we directly tested whether the *k*-mers derived by VirStrain can match commonly seen non-viral reads, including those from human, bacteria, and bacteriophages. In addition, we tested whether there are *k*-mer matches between different viruses. The result is shown in Table 2.

**Table 2:** The number of *k*-mer matches between each type of virus and other two viruses, the human genome, bacteria, and bacteriophages. The human genome, 2,770 complete representative bacterial genomes, and 3,725 complete phage genomes are downloaded from NCBI RefSeq.

| Virus |  | SARS-CoV-2 | H1N1 | HIV | Human | Bacteria | Phage |
| --- | --- | --- | --- | --- | --- | --- | --- |
| name | # $k$-mers | | (HA) | (Gag) | (GRCh38) | | |
| SARS-CoV-2 | 34,754 | - | 0 | 0 | 1 | 0 | 0 |
| H1N1 (HA) | 687,818 | 0 | - | 0 | 13 | 12 | 0 |
| HIV (Gag) | 2,073,196 | 0 | 0 | - | 102 | 63 | 5 |

Table 2 shows that most *k*-mers in the VirStrain database do not match the genomes of other species, indicating that VirStrain is not likely to mistaken other species as viral strains. Our experiments of applying VirStrain to real viral metagenomic data in Section 4.8 further confirmed this.

### 4.4 Detecting a reference strain from simulated reads

In this experiment, we compared VirStrain against Kraken2 [34], KrakenUniq [4], Pathoscope2 [13], Sigma [1], and Centrifuge [17] on detecting one reference strain from the input data. Although there are more taxonomic classification tools for sequence classification, other authors have shown that they cannot achieve satisfactory performance on strain-level composition [6]. Thus, we did not include those tools in our comparison.

For each tool, the reference database is constructed using RNA viral strains. As Sigma is computationally expensive, we were not able to construct its database using all strains. To ensure a fair comparison using the same reference database, we built a smaller, lower-resolution database of 200 stains randomly selected from all strains of the three types of viruses.

For each virus, we randomly picked 100 strains/genes from the 200 reference sequences and simulated short reads from each. Thus, there are 300 datasets for three types of viruses. For each dataset, we used ART [14] to simulate 250 bp error-containing Illumina reads with depth of 100X, average insert size of 600 bp, and standard deviation of 150 bp. We identified strains from these simulated reads with VirStrain and five other tested programs and calculated the accuracy for each program.

The performance comparison of different tools is shown in the left panel of Figure 7A. Gag region of HIV shares relatively low similarity and thus it is easier to distinguish different reference genes. As a result, all tools have high accuracy. As H1N1.HA has very high similarity within the same clades or sub-clades, sequence classification tools that are not specifically designed for distinguishing highly similar genomes have low accuracy. We have similar observations for SARS-CoV-2 too. Across all the three viruses, VirStrain has consistently high accuracy. Tie cases were also checked for all tested tools. Only Centrifuge had tie cases (5/100 for SARS-CoV-2 and 9/100 for H1N1 (HA)) and no tie case was found in the output of all other tested tools. In order to test the performance of VirStrain on all strains,we carried out a benchmark experiment for fast-running tools (see Supplementary Section 2.1). The result shows that VirStrain is able to identify all strains in the database while other tools have lower accuracy (Supplementary Figure S3).

**Figure 7:**
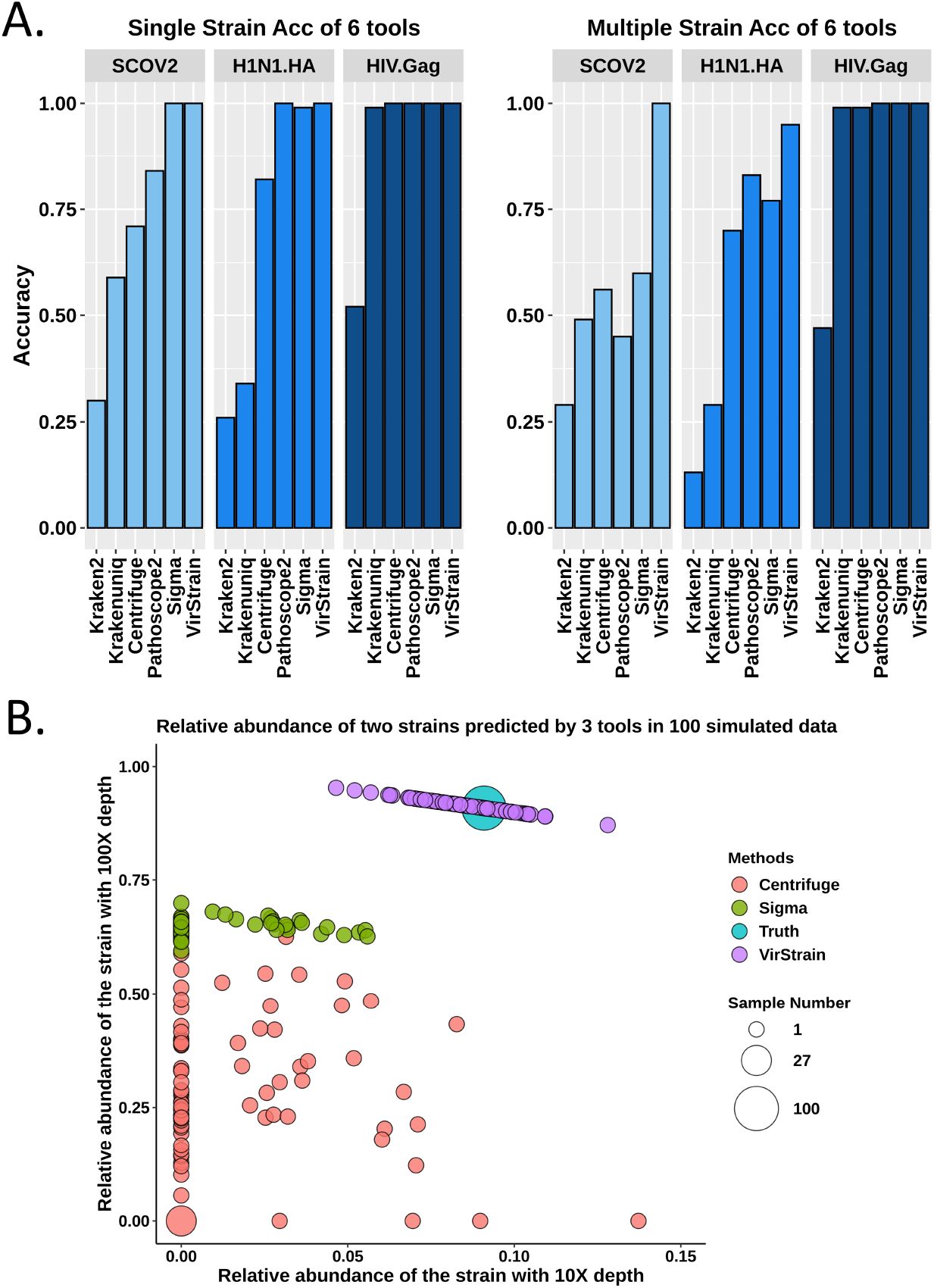
**(A)**. The accuracy (Acc) comparison of 6 tools. There are 100 sets of simulated reads for single-strain datasets and 100 for multi-strain datasets. For each set of multi-strain simulated reads, there are two strains with 100X and 10X coverage, respectively. **(B)**. Predicted relative abundances across 100 simulated SARS-CoV-2 two-strain datasets for VirStrain, Sigma, and Centrifuge. The size of each point (circle) represents the number of simulated data sets. “Truth” refers to the ground truth of the relative abundance of the 2 strains in each data set, which is calculated by normalizing the sequencing depth (100X and 10X).

VirStrain extracted about 30,000 *k*-mers out of roughly 300,000 *k*-mers from the input reference genomes. As shown in Figure 7A, using all *k*-mers in Kraken2 does not render satisfactory accuracy. Based on this, it’s noted that by selecting intelligently chosen unique combinations of k-mers centered around SNVs, strain distinguishment performs as well as if not better than the same program comparing all possible k-mers. Similarly, we observed decreased accuracy if we use all possible *k*-mers in VirStrain. Thus, using selected *k*-mers by the greedy covering algorithm is important to VirStrain.

### 4.5 Detecting multiple strains from simulated data

Multi-strain infection is not rare for RNA viruses, especially the ones with high mutation rates such as HIV. Usually, if one strain dominates the virus population, the minor strains tend to be missed. To mimic this situation, we constructed two-strain datasets that consist of a major strain (100x coverage) and a minor strain (10x coverage). Similar to “single-strain” datasets, we constructed 100 datasets of simulated reads for each type of virus. Each set contains simulated reads from two randomly selected reference sequences. The read simulation process is the same as the single strain experiment. As we know there are two strains, the accuracy of a tool is defined as the ratio of correctly identified strains to the total number of the two most possible strains output by each tool. The performance comparison is shown in the right panel of Figure 7A. Although the accuracy of VirStrain decreases a little for H1N1.HA (from 1.0 to 0.95) compared to the single-strain experiment, it maintains the accuracy of 1.00 for SARS-CoV-2. And it outperforms other tools by about 10% on H1N1(HA) and 38% on SARS-CoV-2.

#### 4.5.1 Relative abundance computation

For identified strains, VirStrain also outputs its sequencing coverage, which can be used to compute relative abundance for multi-strain infection. As Figure 7A shows, accuracy of Kraken2, KrakenUniq, and Pathoscope2 on the SARS-CoV-2 multiple-strain data sets is lower than 0.5. Thus, we didn’t include them in the comparison. Sigma and Centrifuge were able to return the strains’ abundances in the outputs. Therefore it is convenient to calculate the relative abundance for each strain.

Figure 7B shows that the relative abundance estimated by VirStrain is closer to the ground truth than others. Sigma and Centrifuge failed to detect the minor strain in many datasets. Thus, many data points are aligned with x-value 0.00. In addition, they have more variations about the relative abundance computation for different datasets even though the ground truth keeps the same (100x vs 10x).

### 4.6 VirStrain detects the closest relative for novel strains

When a strain is not present in the reference database, VirStrain will output its closest relative in the database. Here, we define the closest relative as the strain in the database that is most similar to the query strain identified by MegaBLAST [38]. In order to test VirStrain on returning the closest relative for novel strains, we created multiple simulated read sets from mutant strain genomes.

In order to test the ability of different tools on detecting the closest relative, we need to reconstruct our reference genome set by choosing *only the sequences that can be correctly identified by all tools*.

Thus, we used 53 SARS-CoV-2 genome sequences that can be identified correctly by all tools in the single-strain experiment. Then, we used simuG program [37] to simulate random point mutations to each of these genomes. According to Dorp et al. [32], the average number of mutations in the SARS-CoV-2 strains is 9.6. Thus, we simulated mutant genomes with 5, 7, 9, 11, 13 random point mutations from the raw genome sequences and marked these newly obtained genomes as M5, M7, M9, M11, M13. In total, there are 265 (53*5) mutant genomes and 53 raw (i.e. reference) genomes. Then, we simulated short reads from these mutant and raw genomes using the same parameters as other experiments. Thus we have 318 (265 mutant and 53 raw) datasets as inputs. For each dataset, as it only contains reads simulated from one strain, we thus only keep the top 1 output by different tools.

Figure 8 shows that VirStrain and Sigma are able to find the correct closest relatives in all data sets, while the other 3 tools failed to output the correct strains in some cases. This is consistent with the experimental results of Sigma [1], which tested this function on multiple datasets.

**Figure 8:**
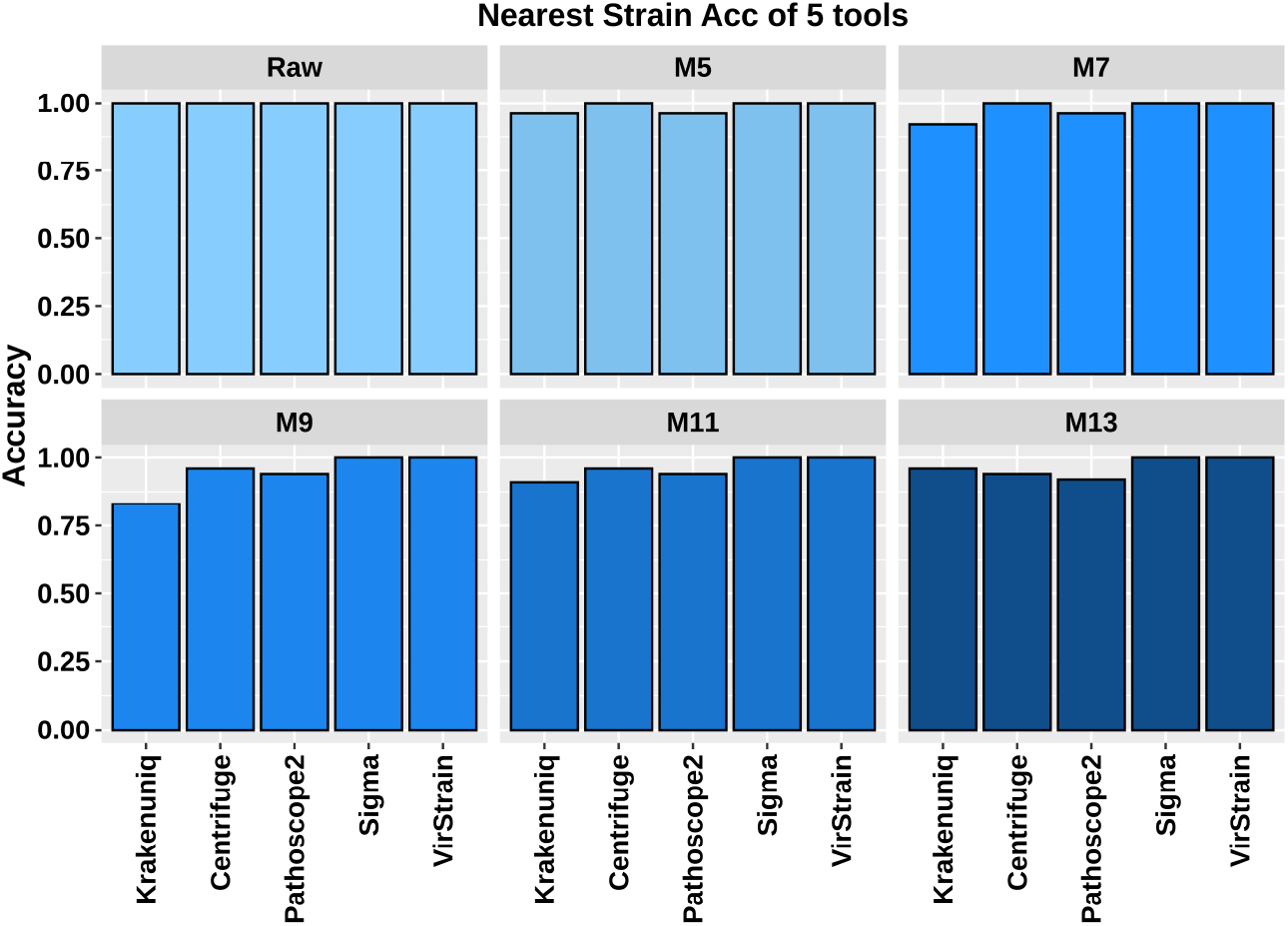
The accuracy comparison of 5 tools on detecting the closest relative in 318 simulated datasets. The 53 strains used in this experiment can be correctly identified by all tools in Figure 7. “Raw” means the dataset from the reference genomes and M5, M7, M9, M11, M13 represent datasets simulated from mutant strains. There are a total of 53 data sets for each group and each dataset contains one strain.

This experiment demonstrated that the performance of our tool is as good as the mapping-based tool Sigma in identifying the closest relative. It is noteworthy that Figure 8 looks better than Figure 7 for several tested tools because this experiment only used 53 strains that are correctly identified by all tools in the single-strain experiment.

To further test the robustness of VirStrain, we applied VirStrain to detect the closest relative in a larger simulated dataset (see Supplementary Section 2.2). The result shows that VirStrain can still identify correct closest relatives in all the 600 simulated datasets (Supplementary Figure S4).

### 4.7 Running time comparison

To evaluate the computational efficiency of VirStrain, we compared the running time of the tested tools on the simulated data and recorded the result in Table 3. One real metagenomic sequencing data (SRR10971381) is also used to compare the computational efficiency due to its large data size. The reference genome of the strain (MN908947) in this dataset (SRR10971381) can be found in the database of all tested tools, so it’s a fair test strain. VirStrain has similar running time to Centrifuge and KrakenUniq but runs significantly faster than Pathoscope2 and Sigma. All the experiments were tested on an HPCC CentOS 6.8 node with 2.4Ghz 14-core Intel Xeon E5-2680v4 CPUs and 128 GB memory. We used 8 threads for all tools.

**Table 3:**
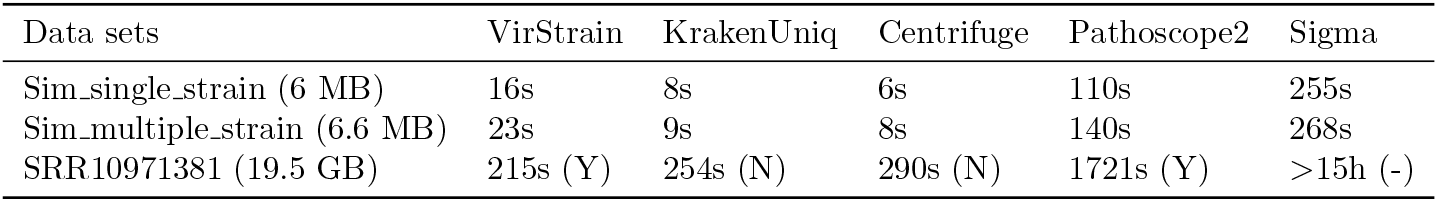
Running time of five tested tools on simulated and real data. Sim_single_strain and Sim_multiple_strain represent simulated single-strain and multiple-strain datasets, respectively. For real data, the identification result is represented by Y and N, where Y means correct identification and N means wrong identification. Sigma doesn’t have the identification result due to its long running time.

These results indicate that VirStrain achieved much higher accuracy than those computationally efficient tools such as KrakenUniq and Centrifuge while maintaining comparable speed. It also outperforms those mapping-based tools like Sigma and Pathoscope2 on both accuracy and speed.

### 4.8 VirStrain detects SARS-CoV-2 strains from real sequencing data

To evaluate the performance of VirStrain in SARS-CoV-2 identification, we conducted experiments on 29 real sequencing datasets (see Table 4), which were sampled from patients of different geographical regions. The samples were sequenced using different platforms such as Illumina, BGI-Seq, and Ion Torrent and may not have complete assemblies available. Viral metagenomic sequencing data are highlighted in dark gray blocks. Out of the 29 samples, 4 samples have their SARS-CoV-2 strains present in the VirStrain database. 16 of them have available complete genomes (the samples marked with red in Table 4). When the complete or near complete viral genomes are given, we used MegaBLAST to find the most similar strain in the VirStrain database and used it as the ground truth. The results are shown in Table 4.

**Table 4:**
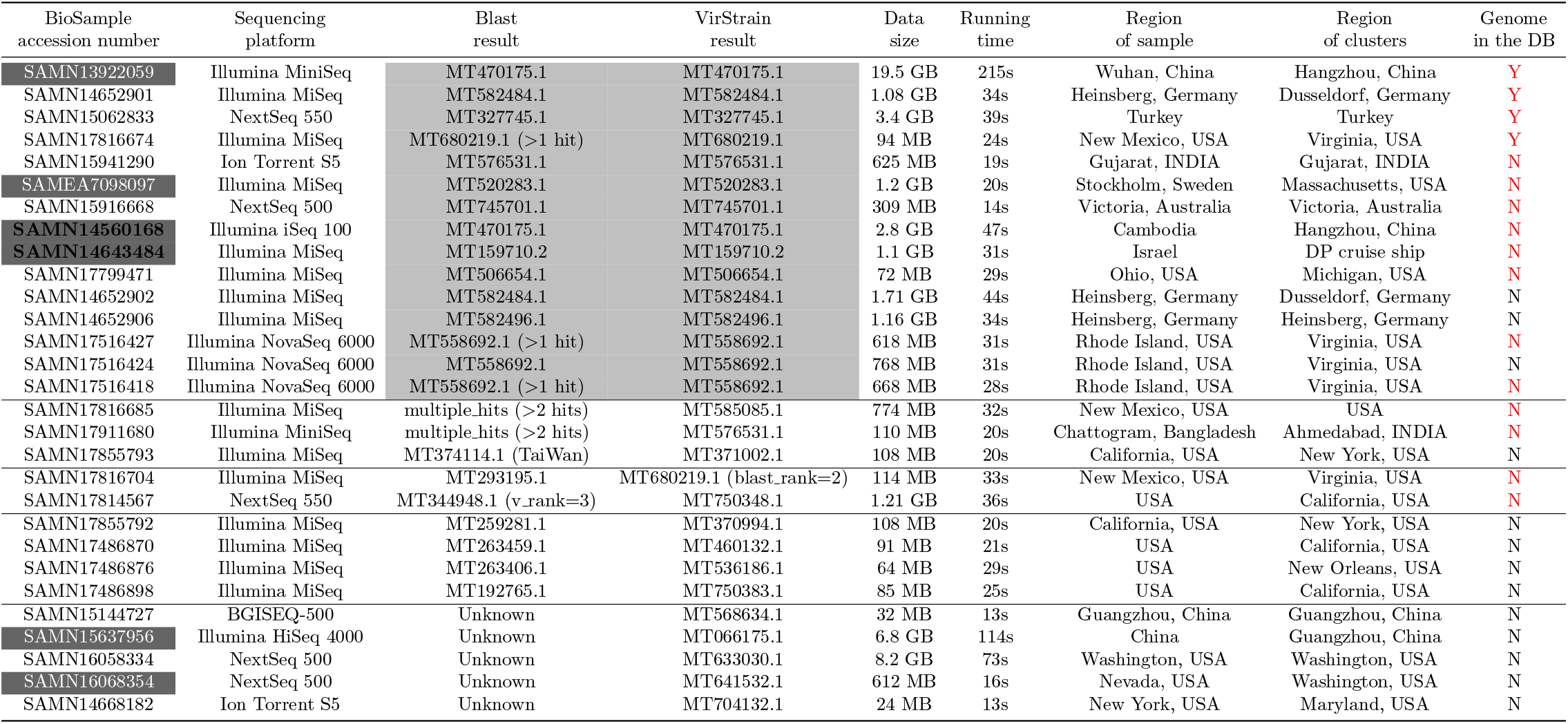
The VirStrain identification result of 29 real sequencing datasets. “Unknown” in the column “Blast result” means that the complete genome of that dataset is not available. “Region of clusters” is the output of VirStrain based on the metadata associated with the reference strains in each cluster. For clusters containing more than one reference strain, we use the majority vote to get the geographical region information. “Genome in the DB” represents whether the complete genome of that dataset can be found in the reference database of VirStrain, yes (Y) or no (N), and the red character means these samples have complete genomes. “DP cruise ship” refers to the Diamond Princess cruise ship. The datasets in the dark gray block means these datasets belong to metagenomic sequencing datasets. “v rank” represents the ranking of the strain in the output of VirStrain. “blast rank” represents the ranking of the strain in the output of Blast.

By comparing the metadata of the output strain by VirStrain and the known information associated with each sample, we can conclude that the derived and known geographical information is generally consistent for all datasets. For most cases where the complete genomes are available, the strains returned by VirStrain are the same as the output of MegaBLAST. There are four cases where VirStrain output different results from MegaBLAST. Of the four cases, MegaBLAST output multiple hits for two. For the other two, the strain identified by VirStrain are very close to MegaBLAST. As VirStrain uses short reads as input, this indicates that its accuracy is comparable to highly accurate alignment tools that take genomes as input.

The first sample SAMN13922059 is actually from a patient in Wuhan, China, whose sample was used to generate the first reference genome of SARS-CoV-2 [35]. In the output of VirStrain this first reference genome is located in a cluster with other 47 SARS-CoV-2 strains, which all belong to clade 19A defined by nextstrain. In this cluster, there are two main geographical locations: Wuhan and Hangzhou, China. As Hangzhou’s cases are slightly more than Wuhan, we used Hangzhou in column “Region of clusters”. This is one current limitations of VirStrain. These 48 strains cannot be divided into single-strain clusters.

There are 2 very interesting samples in Table 4: SAMN14560168 and SAMN14643484 (bold font). SAMN14560168 is from the first COVID-19 patient of Cambodia, who had been to China before being admitted to the hospital. The identification result of VirStrain shows that its closest relative is MT470175.1, which is from China. Thus, the result indicates that this Cambodia patient could be infected in China, which is consistent with this patient’s travel history. Another interesting case, SAMN14643484, is from Israel and its closest relative identified by VirStrain is from the Diamond Princess cruise ship. According to the sample information at NCBI, this patent was indeed a passenger of the cruise ship and got infected by SARS-CoV-2 there.

These results show that that VirStrain is able to identify SARS-CoV-2 strains from real sequencing data with or without assembled genomes. In addition, VirStrain also provides information that can be very useful for tracking the virus spread.

### 4.9 VirStrain identifies 5 strains from HIV mock data

In this experiment, we applied VirStrain to a mock dataset (SRR961514) containing real sequencing data from five HIV strains. The authors mixed five HIV strains (JRCSF, 89.6, NL43, YU2, and HXB2) and conducted Illumina sequencing [8]. Using the reads as input, VirStrain can detect 5 strains from its reference database and predict their sequencing depth. Based on the predicted sequencing depth, we calculated the relative abundance by normalizing the depth of each identified strain.

To compare the predicted abundance with the ground truth, we applied the chi-square test and got the p-value 0.9998, which indicates that the distribution of the predicted abundance by VirStrain is not statistically different from the ground truth (Figure 9). This experiment demonstrates the ability of VirStrain in identifying multiple strains in one sample.

**Figure 9:**
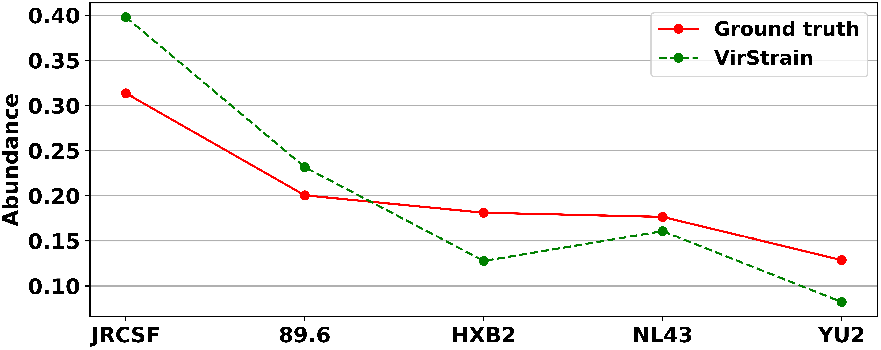
Abundance comparison of HIV mock data between the ground truth and VirStrain. The true average abundances sorted in descending order are: 31.35%, 20.04%, 18.11%, 17.65%, 12.86%

## 5 Discussion

Our large-scale benchmark experiments against several other popular strain-level analysis tools demonstrated the high accuracy of VirStrain on detecting reference strains from short reads. But there are still cases where VirStrain cannot return the exact strain. One limitation of current method is the ambiguity of detecting low abundance viral strains (depth *<* 10x). As mentioned in Section 4.3, there could be multiple best matches when the depth is smaller than 10X. As not all SNV sites can be covered by the reads, strains of high similarity and with a large number of shared SNVs can form a tie case with the same score. With coverage bigger than 10x, the tie cases become very rare and the top 1 strain identified by VirStrain is the correct strain in the sample. This limitation caused by low coverage and high similarity is also observed in other tested tools. Instead of outputting a wrong strain, VirStrain outputs all with the correct strain being one of them, which can inform the users of this ambiguity. It is our future work to design more accurate algorithms for addressing this limitation.

In the case of detecting multiple strains in one sample, there is a tradeoff between resolution and accuracy. Specifically, if there are multiple strains sharing a large number of SNV sites, they will be clustered in the list ranked by *V score*, which can pose false positive detection when one of the strain exists in the underlying sample. Thus, we do not reuse the SNV sites so that the output strains are representative ones in a sample rather than near duplicate ones. Essentially, the iterative search procedure poses a constraint on the number of different SNVs between strains in the same sample. Strains with too few differences will be missed by VirStrain. In order to provide the guidance on the number of expected different SNVs, we tested a hard case for our method. The input data contains reads from three strains, with one being the major one (100x) and other two being minor ones (10x). In addition, two of the three strains are highly similar with less than 10 different SNV while the rest one has more than 3 different SNVs. The result can be found in Supplementary Section 2.5. It shows that VirStrain may miss the low abundance strain that differ by less than 10 SNV sites for SARS-CoV-2 in the multi-strain infection cases.

Currently we derive *k*-mers from aligned reference genomes. We noticed some alignment errors especially at sites with consecutive insertions or deletions. As a result, we tend to exclude columns with many indels, which may lead to clusters containing multiple genomes in the end. Ideally, we want to derive *k*-mer sets for reference genomes without relying on alignment programs, which is our future work.

## 6 Conclusions

In this work, we implemented a strain identification tool for short reads. We designed a greedy covering algorithm to divide reference genomes into multiple clusters so that the genomes in each cluster possess unique set of *k*-mers.

VirStrain shows higher accuracy than other tested tools across all benchmark datasets with different complexity. One of VirStrain’s advantages is that it can detect the closest relative for novel strains. For example, VirStrain identified the most similar strain from datasets containing mutant genomes that are not in the database and demonstrated the same accuracy as using MegaBLAST [38] on complete genomes. Another advantage of VirStrain is to detect multi-strain infection cases. We demonstrated this by using VirStrain on both simulated and real sequencing data.

We also plan to adapt this method for bacterial strain analysis, which is an important component in precise diagnosis and treatment for clinical data. Some bacteria already have a large number of sequenced strains, which can benefit from our fast *k*-mer set derivation method.

## 7 DATA AVAILABILITY

All data and codes used for this study are available online or upon request to the authors. The source code of VirStrain is freely available at https://github.com/liaoherui/VirStrain.

## 8 ACKNOWLEDGEMENTS

This work was supported by Hong Kong RGC GRF 9042828 and HKIDS (9360163).

