## Supplementary Data for "VirStrain: a strain identification tool for RNA viruses"

March 2021

### 1 Supplementary Methods

#### 1.1 *k*-mer extraction from SNV sites

We will extract $k$-mers from these SNV sites, with the center base of each $k$-mer coming from this site. The complete $k$-mer is then constructed by including $\frac{k-1}{2}$ bases to the left and right side of a variation site. Supplementary Figure S1 shows an example of $k$-mer extraction.

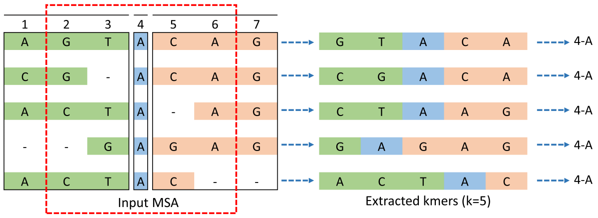


**Supplementary Figure S1.** The process of $k$-mer extraction. Given the chosen site, $k$-mers are extracted from the centering base and neighbouring sequences.

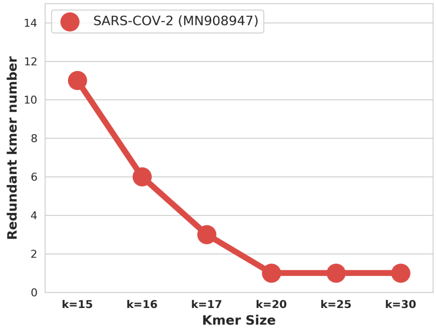


**Supplementary Figure S2.** The relationship between $k$-mer size and the number of redundant $k$-mers. When $k \geq 20$, the only repetitive $k$-mer is a repeat of “A”, which is not used in our method.

If $k$-mers repeat at different sites in the reference genomes, false positive strain identification will happen. Intuitively, bigger $k$-mers tend to appear less frequently than small $k$-mers. We thus investigate how many times we will observe the same *k*-mer at different sites in an MSA. As the reference genomes SARS-CoV-2 have high sequence similarity and larger genome sizes than other known RNA viruses, we conducted this experiment on SARS-CoV-2. The results are summarized in Supplementary Figure S2. As the figure shows, with the increase of *k*, the repeat numbers of the *k*-mer at different sites reduce quickly. Thus, either large *k* can be chosen or we can directly exclude those columns that contain repetitive *k*-mers in order to fulfill the assumption.

### 2 Supplementary Experiments

#### 2.1 Benchmark experiment on all strains

In order to test the performance of VirStrain on all strains, we carried out a benchmark experiment with VirStrain, KrakenUniq, and Centrifuge. As Sigma and Pathoscope2 are more computationally expensive than other tools, they are not compared in this experiment. Kraken2 is also not compared due to its low accuracy on the single-strain dataset.

In this experiment, one of the SARS-CoV-2 strains in the reference database was randomly selected as the test strain. We simulated reads from it and used the reads as input to the three tools. This process is repeated for all other strains. The accuracy is computed for all the strains and shown in Supplementary Figure S3. VirStrain is able to identify all strains in the database while other tools have lower accuracy.

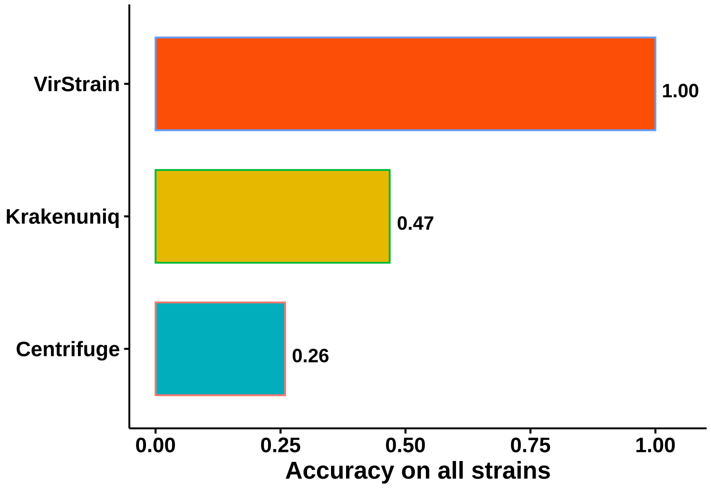


**Supplementary Figure S3.** The accuracy comparison of 3 tools on all strains in the database. These three tools have fast speed and also reasonable accuracy in the small scale experiment on 200 reference genomes.

#### 2.2 A large-scale experiment of novel strain detection

To further test the robustness of VirStrain, we applied VirStrain to detect the closest relative in a larger simulated dataset. Sigma was not included in this experiment because of its long running time. We randomly selected 100 strains from all the reference genomes. Then we generated 500 mutant strains from these 100 strains with 5, 7, 9, 11, 13 random point mutations. With ART, we simulated 600 datasets (for 500 mutant strains and 100 reference strains) and used them as input to VirStrain. VirStrain can still identify correct closest relatives in all the 600 simulated datasets (Supplementary Figure S4).

#### 2.3 VirStrain detects possible minor strains from Washington State datasets

For RNA viruses with fast replication rates, it is not rare to see multiple strains infecting the same host. For COVID-19, is it possible to see more than one strain (or haplotype) in the same host? Reconstructing the complete haplotype from short reads is an active research area itself and is beyond the scope of this work. But we can examine whether a different local haplotype segment exist in a sample. If we find one such local haplotype segment, we can add it to

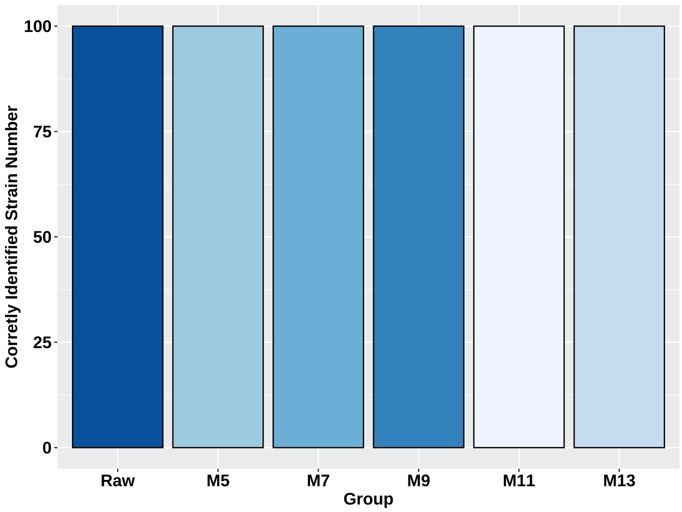


**Supplementary Figure S4.** The performance of VirStrain on detecting the closest relative on 600 simulated datasets. "Raw" means the original dataset and M5, M7, M9, M11, M13 represents datasets simulated from mutant strains. Each dataset contains one strain.

our reference sequence database and search for it in other samples using VirStrain. Being able to detect the same local haplotype in multiple samples provides strong evidence behind the presence of a new haplotype.

Towards this goal, we first assembled reads from sequenced SARS-CoV-2 samples at NCBI SRA using Megahit [4] and then mapped reads against the assembled genome. We found some relatively high-frequency mutations in a Washington State sample (SAMN15678406) from USA (Supplementary Figure S5). Based on the results of the reads mapping and a statistical model that considers the base quality, sequencing errors, coverage, and the number of reads covering the same SNV events, we obtained a possible local region of a minor strain, which we named as k59_23. We used nextstrain [2] to annotate k59_23 and found that there is a mutation G25358C that will cause the amino acid mutation D1259H of the surface protein. The statistical model used to find the possible local haplotype is described below.

There are four observed single nucleotide variants (SNVs) with frequencies much higher than sequencing errors at different loci of reference genome based on the read mapping results: 25345-A, 25355-C, 25357-C, 25358-C. We regard other types of SNVs at these four loci as sequencing errors because their frequencies are very low. We are not aiming to reconstruct the complete (genome-scale) haplotype, which is an active research field itself and is beyond the scope of this work. Instead, we will construct a local haplotype region containing SNV sites that are close enough to be covered by the same reads.

In the reference genome, the bases at the four loci are 25345-C, 25355-G, 25357-A, 25358-G. Thus, of the $2^4 - 1 = 15$ combinations of bases (excluding the reference one), our goal is to determine the most possible combination, which can define a local haplotype. For the sake of convenience, we use $k = 1, 2, 3, 4$ to denote these four loci. And we define the following notations: **a)** $q_k^{(j)}$ is the Phred quality score of the base in read $j$ when this base mismatches the base of reference genome at locus $k$ in the alignment between the read $j$ and the reference genome. If the base in read $j$ matches the base of reference genome at locus $k$ in the alignment, or read $j$ does not cover locus $k$ in the alignment, the value of $q_k^{(j)}$ will be set to 0. **b)** $c_k^{(j)} \in \{0, 1\}$ is an indication function. $c_k^{(j)} = 1$ if the base in read $j$ is not a sequencing error at locus $k$ in the alignment. $c_k^{(j)} = 0$ if the base in read $j$ is a sequencing error at locus $k$ in the alignment, or read $j$ does not cover locus $k$ in the alignmen. **c)** $m_k$ is the number of reads whose bases match the SNV at locus $k$ in the alignments between the reads and the reference genome. **d)** $d_k$ is the number of aligned reads that are aligned to locus $k$. **e)** $p_k^{(j)}$ is the probability that read $j$ supports the SNV at locus $k$. And We use the equations(1) to calculate $p_k^{(j)}$.

$$p_k^{(j)} = \begin{cases} 1 - 10^{-\frac{q_k^{(j)}}{10}} & \text{if } q_k^{(j)} \neq 0 \; and \; c_k^{(j)} = 1 \\ 0 & \text{if } q_k^{(j)} = 0 \; and \; c_k^{(j)} = 1 \\ 10^{-\frac{q_k^{(j)}}{10}} \times \frac{m_k}{d_k} & \text{if } q_k^{(j)} \neq 0 \; and \; c_k^{(j)} = 0 \\ \frac{m_k}{d_k} & \text{if } q_k^{(j)} = 0 \; and \; c_k^{(j)} = 0 \end{cases} \tag{1}$$

There are four cases in Equation (1), also as shown in Figure S1. **In case 1**, the base in read $j$ aligned to locus $k$ of the reference genome matches the SNV at locus $k$. In this case we can use the Phred quality score to calculate the SNV probability directly. **In case 2**, the base in read $j$ aligned to locus $k$ of the reference genome matches the base in the reference genome and thus this read does not support the SNV event. So it is reasonable to assume that the probability of read $j$ containing SNV at locus $k$ is 0. **In case 3**, the base in read $j$ aligned to locus $k$ of the reference genome is a sequencing error. The true base can be the reference one or the SNV. Thus, we can use the error rate and the nucleotide frequencies observed at the locus $k$, to estimate the probability of read $j$ supporting this SNV event. **In case 4**, if the alignment between read $j$ and the reference genome does not cover locus $k$ (missing information), we use the nucleotide frequencies observed at the locus $k$ to estimate the probability of read $j$ supporting this SNV event.

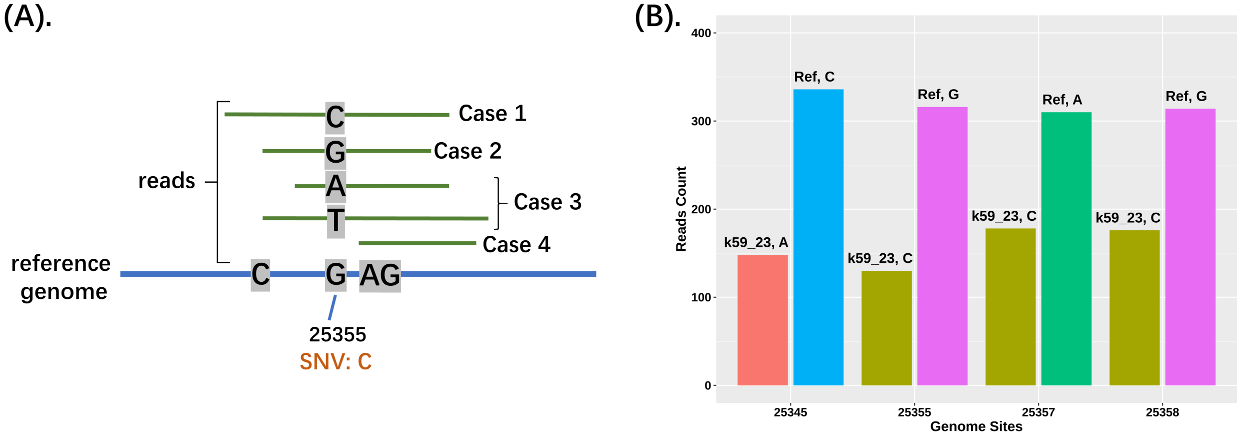


**Supplementary Figure S5.** A. Four cases for equation (1) at locus 25355. B. The read count comparison matching the reference and k59_23 at the four SNV sites, respectively.

Similarly, we define $r_k^{(j)}$ as the "reference probability" that read $j$ supports the reference base at locus $k$. And we can use the equation (2) to calculate $r_k^{(j)}$.

$$r_k^{(j)} = \begin{cases} 10^{-\frac{q_k^{(j)}}{10}} & \text{if } q_k^{(j)} \neq 0 \; and \; c_k^{(j)} = 1 \\ 1 & \text{if } q_k^{(j)} = 0 \; and \; c_k^{(j)} = 1 \\ 10^{-\frac{q_k^{(j)}}{10}} \times (1 - \frac{m_k}{d_k}) & \text{if } q_k^{(j)} \neq 0 \; and \; c_k^{(j)} = 0 \\ 1 - \frac{m_k}{d_k} & \text{if } q_k^{(j)} = 0 \; and \; c_k^{(j)} = 0 \end{cases} \tag{2}$$

Based on the SNV probability equation (1) and reference probability equation (2), we define Equation (3) to calculate the probability that read $j$ is consistent with haplotype $i$, where $i$ is one of the 15 SNV combinations.

$$Pr(H^{(i)}|R^{(j)}) = \prod_{k=1:4, h_k^{(i)}=1} h_k^{(i)} p_k^{(j)} \times \prod_{k=1:4, h_k^{(i)}=0} (1 - h_k^{(i)}) r_k^{(j)} \tag{3}$$

$R^{(j)}$ is the read $j$. $H^{(i)} = \{h_k^{(i)} | k = 1, ..., 4\}$ is the haplotype $i$. $h_k^{(i)} \in \{0, 1\}$ is an indication function such that $h_k^{(i)} = 1$ if haplotype $i$ contains the SNV at locus $k$ and $h_k^{(i)} = 0$ if the haplotype $i$ contains the reference base at locus $k$ (e.g. the indication function for haplotype AGCG will be $\{1, 0, 1, 0\}$). To find out the most possible haplotype that is different from the reference genome, we use Equation (4) to calculate the normalized weight of each haplotype based on the read mapping quality of a set of given reads. This weight denotes the consistency between each haplotype and the set of reads.

$$Weight(H^{(i)}|R) = \frac{\sum\limits_{R^{(j)} \in R} (1 - 10^{-\frac{Q^{(j)}_{map}}{10}}) Pr(H^{(i)}|R^{(j)})}{\sum\limits_{R^{(j)} \in R} (1 - 10^{-\frac{Q^{(j)}_{map}}{10}})} \tag{4}$$

where $Q^{(j)}_{map}$ is the mapping quality of read $j$ obtained from bowtie2 [3]. $(1 - 10^{-\frac{Q^{(j)}_{m}ap}{10}})$ denotes the estimated probability that the alignment of read $j$ corresponds to the true region. $R$ is a set of reads that cover and match at least one of the four loci of the reference genome we mentioned above. Table S1 shows the weights of different haplotypes.

We can see that the haplotype **ACCC** at four loci is the most possible one of the fifteen possible haplotypes. Its weight is about $0.2235$, which is about the twice of the second-highest weight. Thus we add this possibly novel haplotype into the reference database and used VirStrain to search for this haplotype in other Washington State samples.

**Supplementary Table S1.** Weights of different haplotypes. The character with blod face is a SNV.

| Haplotype | Weight | Haplotype | Weight | Haplotype | Weight |
|---|---|---|---|---|---|
| **ACCC** | 0.2235 | **ACAC** | 0.0799 | CGA**C** | 0.0321 |
| CCCC | 0.1258 | **A**GAG | 0.0499 | **AC**AG | 0.0284 |
| **A**G**CC** | 0.1160 | **A**GA**C** | 0.0497 | C**C**A**C** | 0.0236 |
| CG**CC** | 0.0816 | **ACC**G | 0.0383 | C**CC**G | 0.0200 |
| **A**G**C**G | 0.0815 | CG**C**G | 0.0323 | C**C**AG | 0.0160 |

In order to validate this finding, we checked whether other samples also contain k59_23. Towards this goal, we added this strain to the VirStrain database and then used VirStrain to detect it in other SRA datasets from Washington State. The output shows that k59_23 was also detected in three other Washington State samples.

The VirStrain detection results of these four data sets are summarized in Supplementary Table S2. Both the statistical analysis and the VirStrain outputs show that a minor strain likely exists in these samples. This indicates the possible utility of VirStrain in detecting multi-infection cases.

| BioSample accession number | VirStrain Result 1 | Relative abundance 1 | VirStrain Result 2 | Relative abundance 2 |
|---|---|---|---|---|
| SAMN15678406 | MT641500.1 | 0.8523 | k59_23 | 0.1477 |
| SAMN15678405 | MT375468.1 | 0.9042 | k59_23 | 0.0958 |
| SAMN15678404 | MT632835.1 | 0.8797 | k59_23 | 0.1203 |
| SAMN15678403 | MT628075.1 | 0.9006 | k59_23 | 0.0994 |

**Supplementary Table S2.** The identification results of VirStrain of four Washington State samples. VirStrain can identify two strains in these data sets. The numbers "1" and "2" represent these 2 strains. The major strains in these four samples returned by VirStrain are different but all have geographical locations from Washington State. The relative abundance of each strain is calculated by normalizing the depth of all identified strains.

#### 2.4 VirStrain predicts HIV subtype and H1N1 clade

The identification result generated by VirStrain can also be used to extract the subtype or clade information of identified strains. Thus, users can obtain the strain subtype or clade information from reads directly without assembly, which is useful for datasets that cannot form quality assemblies. In this experiment, we applied VirStrain to four real datasets and output the subtype or clade information of identified strains using a bottom-to-up method. To know the subtype or clade type of the input dataset, we can check the metadata of "the most possible strain" output by VirStrain. Then, we use the subtype or clade label of the identified strain as the final output.

Supplementary Table S3 shows that the subtype or clade identified by VirStrain is the same as the truth. Besides, the region information is also consistent with the source region of these samples.

| BioSample accession number | Virus type | Subtype/Clade (Truth) | Subtype/Clade (VirStrain) | Region of sample | Major region of cluster |
|---|---|---|---|---|---|
| SAMN10225085 | HIV | F1 | F1 | Brazil | Brazil |
| SAMN08101778 | HIV | B | B | Brazil | Brazil |
| SAMN01095441 | H1N1 | 1A.3.3.2 | 1A.3.3.2 | Hong Kong | Hong Kong |
| SAMN02719578 | H1N1 | Other-Human | Other-Human | Unknown | Russia |

**Supplementary Table S3.** The VirStrain clade/Subtype prediction result of 4 real sequencing data. "Subtype": the subtype of HIV strain is obtained according to literature [1]. "Clade": the clade of H1N1 strain is estimated by the Swine H1 clade classification tool of Influenza Research Database (https://www.fludb.org).

#### 2.5 Limitation posed by iterative search

As many reference strains share high sequence similarity, after finding the best match in the reference genomes, VirStrain will remove the matched SNV sites before the next iteration so that other possible strains with different SNV sites can achieve better $Vscore$. Without removing the matched SNV sites, strains that share high similarity with the identified strain will naturally returned as other possible strains, incurring false positive identifications. However, there is a tradeoff between the false positive identification and sensitivity. When the SNV sites of the best matched strain are removed, the strains that are highly similar to the best matched one may be missed by VirStrain. To investigate this, we constructed a difficult case where three strains with different similarities co-exist in one sample and tested the performance of VirStrain.

Thus, the iterative search procedure poses a constraint on the number of different SNVs between strains in the same sample. Strains with too few differences will be missed by VirStrain. In order to provide the guidance on the number of expected different SNVs, we tested a hard case for our method. The input data contain reads from three strains, with one being the major one (100x) and other two being minor ones (10x). In addition, two of the three strains are highly similar with only 1 different SNV while the rest one has more than 3 different SNVs (strains with 3, 5 and at least 10 different SNVs were used and considered as 3 groups in this experiment). In total, there are 30 simulated datasets and 10 simulated datasets in each group. Supplementary Table S4 shows VirStrain is always able to identify strains with high abundance and more than 3 different SNVs but misses the strain with 1 different SNV. In addition, we found the median number of best matches decrease with the increase of the number of different SNVs. Based on these results, we recommend users to consider 10 as a minimum number to identify other possible strains in multi-strain infection cases. For low abundance strains identified in all tie cases, VirStrain will output multiple best matches in descending order of $Vscore$ of the first iteration. We found the correct strain was always the top 1 strain under this sorting rule, which could be a useful information for users to identify the correct strain among multiple best matches.

| Group | Depth | Identification result | Tie cases | Median number of best matches |
|---|---|---|---|---|
| (H, 1, 3) | (100X, 10X, 10X) | (10, 0, 10) | (0, 0, 4) | (1, 0, 6) |
| (H, 1, 5) | (100X, 10X, 10X) | (10, 0, 10) | (0, 0, 4) | (1, 0, 5) |
| (H, 1, >=10) | (100X, 10X, 10X) | (10, 0, 10) | (0, 0, 4) | (1, 0, 2) |

**Supplementary Table S4.** The performance of VirStrain on 30 3-strain simulted datasets with different settings (10 simulated datasets in each group). "H" in "Group" column represents the high abundance strain, and other two values in the tuple represent the strain with x SNVs (x refers to the value in the tuple). For "Identification result", "Tie cases", "Median number of best matches" columns, each value in the tuple represents the count of these attributes for correspond strain among all simulated datasets of that group. For example, "(10, 0, 10)" in the first row, "Identification result" column represents high abundance strain and the strain with 3 different SNVs can be identified in all 10 datasets while the strain with 1 different SNV is lost in all tested datasets.

### 3 Supplementary Figures

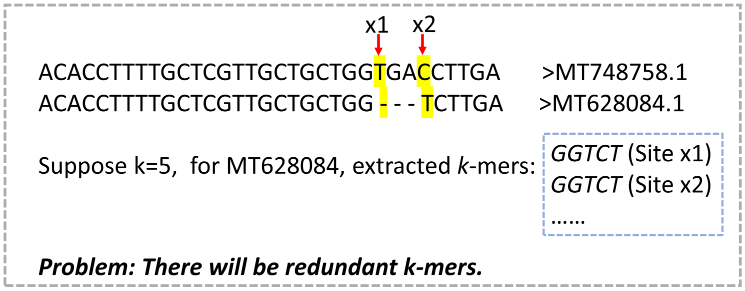


**Supplementary Figure S6.** An example showing the problem caused by dash.

A.

| Strain ID | Cls info | SubCls info | Vscore | Valid map rate | Site coverage | Strain depth | Strain info | SNV freq |
|---|---|---|---|---|---|---|---|---|
| >> Most possible strains | | | | | | | | |
| >MT451123.1 | Cluster2830_2 | NA | 1 | 2877/2877 | 2877/2877 | 73.62872975 | Name:AUS/VIC223/2020\|Clade:19A | {'574-G': 97, '592-G': 57,...} |
| >> Other possible strains | | | | | | | | |
| Can not detect other strains. | | | | | | | | |
| >> Highest_Map_Strains (Could be FP): | | | | | | | | |
| >MT451123.1 | Cluster2830_2 | NA | 1 | 2877/2877 | 2877/2877 | 73.62872975 | Name:AUS/VIC223/2020\|Clade:19A | {'574-G': 97, '592-G': 57,...} |
| >> Top10_Score_Strains: | | | | | | | | |
| >MT451123.1 | Cluster2830_2 | NA | 1 | 2877/2877 | 2877/2877 | NA | Name:AUS/VIC223/2020\|Clade:19A | NA |
| >MT641682.1 | Cluster950_2 | Not_record | 0.9996053 | 2876/2877 | 2876/2877 | NA | Name:AUS/VIC509/2020\|Clade:19A | NA |
| >MT451218.1 | Cluster2887_3 | Not_record | 0.999361 | 2875/2877 | 2875/2877 | NA | Name:AUS/VIC326/2020\|Clade:19A | NA |
| >MT451197.1 | Cluster2879_1 | Not_record | 0.9990321 | 2874/2876 | 2874/2876 | NA | Name:AUS/VIC305/2020\|Clade:19A | NA |
| >MT633044.1 | Cluster1615_3 | Not_record | 0.9989005 | 2874/2877 | 2874/2877 | NA | Name:USA/WA-S413/2020\|Clade:19A | NA |
| >MT451079.1 | Cluster2812_1 | Not_record | 0.9988911 | 2874/2876 | 2874/2876 | NA | Name:AUS/VIC176/2020\|Clade:19A | NA |
| >MT407654.1 | Cluster2152_2 | Not_record | 0.9986045 | 2873/2877 | 2873/2877 | NA | Name:CHN/OS1/2020\|Clade:19A | NA |
| >MT459850.1 | Cluster2619_2 | Not_record | 0.9985481 | 2873/2877 | 2873/2877 | NA | Name:GRC/134_32124/2020 \|Clade:19A | NA |
| >MT358709.1 | Cluster3485_2 | Not_record | 0.9985481 | 2873/2877 | 2873/2877 | NA | Name:USA/ID-UW-3892/2020\|Clade:19A | NA |
| >MT451220.1 | Cluster2888_2 | Not_record | 0.9985434 | 2873/2877 | 2873/2877 | NA | Name:AUS/VIC328/2020\|Clade:19A | NA |

B.

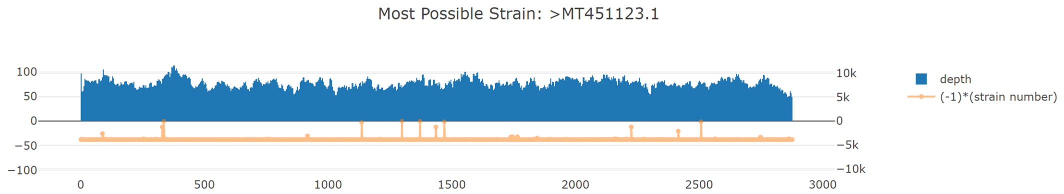


**Supplementary Figure S7.** A. The VirStrain text report of one SARS-CoV-2 simulated dataset (Truth: MT451123.1). The detailed explanation of the columns can be found at the Github website of VirStrain. B. The VirStrain html report of one SARS-CoV-2 simulated dataset (Truth: MT451123.1). X-axis refers to all selected sites by the greedy covering algorithm. The top Y-axis represents the predicted sequencing depth of each site. The bottom Y-axis represents the number of strains sharing this site, which reflects the site uniqueness.

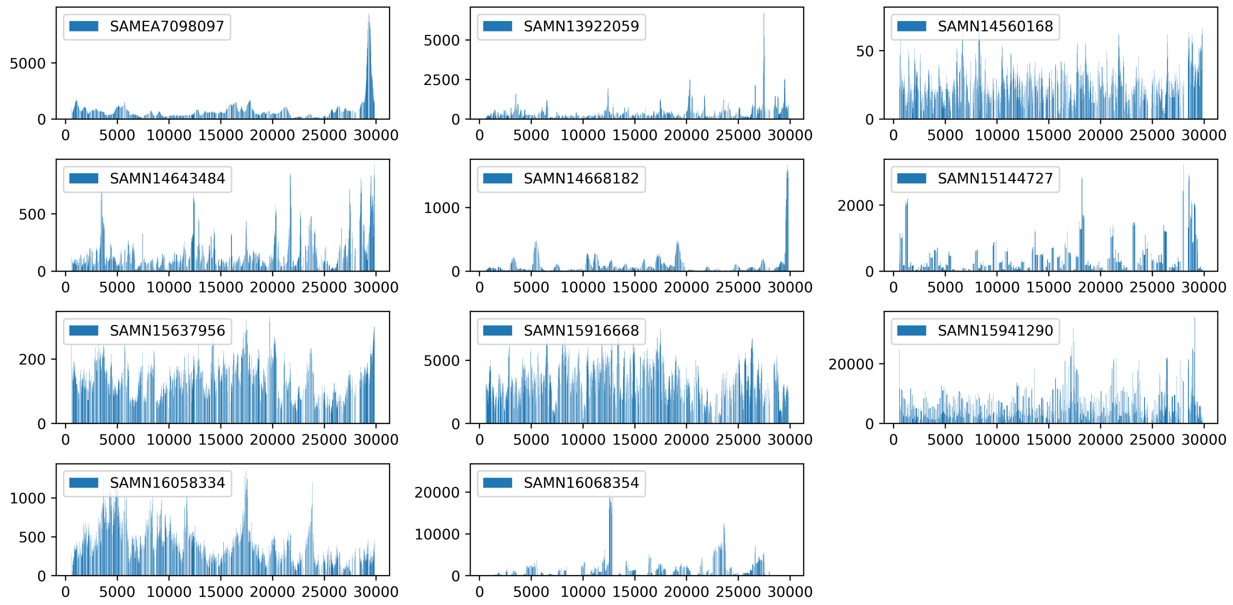


**Supplementary Figure S8.** The sequencing depth distribution of the SNV sites of SARS-CoV-2 genomes in 11 real sequencing datasets. X-axis represents the position of identified genomes. Y-axis represents the sequencing depth of chosen SNV sites by the greedy covering algorithm. The strains shown in these figures are all correctly identified.
